# Synchronous activation of striatal cholinergic interneurons induces local serotonin release

**DOI:** 10.1101/2024.11.03.621726

**Authors:** Lior Matityahu, Zachary B. Hobel, Noa Berkowitz, Jeffrey M. Malgady, Naomi Gilin, Joshua L. Plotkin, Joshua A. Goldberg

## Abstract

Striatal cholinergic interneurons (CINs) activate nicotinic acetylcholine receptors on dopamine axons to extend the range of dopamine release. Here we show that synchronous activation of CINs induces and extends the range of local serotonin release via a similar mechanism. This process is exaggerated in the hypercholinergic striatum of a mouse model of OCD-like behavior, implicating CINs as critical regulators of serotonin levels in the healthy and pathological striatum.

## Main Text

As the main input nucleus of the basal ganglia, the dorsal striatum guides goal directed and habit learning by plastically adjusting how it processes cortical inputs^1,2^. Most of the relevant processes are shaped, at least in part, by the monoamine serotonin (5-HT) – this includes long term depression at corticostriatal synapses^3^, lateral inhibition between spiny projection neurons^4^ and release of another monoamine (dopamine) from nigrostriatal projections^5^. As such, regulation of 5-HT signaling is crucial to striatal function^6^. It has become apparent that monoamines and acetylcholine can intimately shape each other’s release in the striatum: for example dopamine regulates the excitability of striatal cholinergic interneurons^7^ (CINs) and acetylcholine (ACh) release from CINs directly facilitates dopamine release by engaging nicotinic ACh receptors (nAChRs) on nigrostriatal axons^8–12^. Whether such a direct interplay extends to 5-HT is unclear: it is well-established that 5-HT directly regulates the activity of CINs^13,14^, and thus ACh release, but evidence for direct functional control of 5-HT release by ACh is limited^15,16^. The source of striatal 5-HT is a long range projection from the dorsal raphe nucleus^17^, and serotonergic axons within the striatum express nAChRs^15,16^. We therefore sought to determine if CIN-derived ACh can directly facilitate local 5-HT release, in a nAChR-dependent way.

To this end, we stereotaxically inoculated the dorsolateral striatum (DLS) of mice with an adeno-associated virus (AAV) harboring the genetically-encoded 5-HT sensor, GRAB-5HT^18^, and when the sensor was expressed, we prepared acute brain slices through the dorsal striatum (**Fig. 1a**). Intrastriatal electrical stimulation induced a robust increase in GRAB-5-HT fluorescence, which decayed over 10s of seconds, as measured using two-photon laser scanning microscopy (2PLSM) (**Fig. 1b**) (see Methods). Application of the non-selective nAChR antagonist mecamylamine (10 µM) significantly reduced the amplitude of the striatal GRAB-5HT response (averaged over the entire field-of-view; **Fig. 1a-c**), suggesting that, as is the case for dopamine (DA) ^8–11,19,20^, nAChR activation also facilitates striatal 5-HT release [*N = 3* mice, *n = 9* slices, *t*_*16*_ *= 7*.*8, P = 4*.*3*·*10*^*–5*^, linear-mixed effect model (LMEM)]. Bath application of DA (10 µM) had no effect on GRAB-5-HT fluorescence in slices (**Supplementary Fig. 1**), confirming that changes in sensor fluorescence were not the result of nonselective detection of DA^18^. We further confirmed the specificity of GRAB-5-HT fluorescence in our experiments by demonstrating that 1) RS 23597-190 (35 µM; a selective antagonist of the 5-HT_2C_ receptor, on which the GRAB-5HT design is based) eliminated evoked fluorescence (*N = 3* mice, *n = 6* slices, *t*_*10*_ *=* –*4*.*033, P = 0*.*002*, LMEM) (**Supplementary Fig. 2b**,**c**) and 2) citalopram (30 µM; a selective antagonist of 5-HT reuptake) significantly slowed the temporal dynamics of evoked fluorescence (*N = 4* mice, *n = 12* slices, *t*_*22*_ *= –4*.*3, P = 2*.*9*·*10*^*–4*^, LMEM) (**Supplementary Fig. 2d**,**e**).

**Figure 1.**
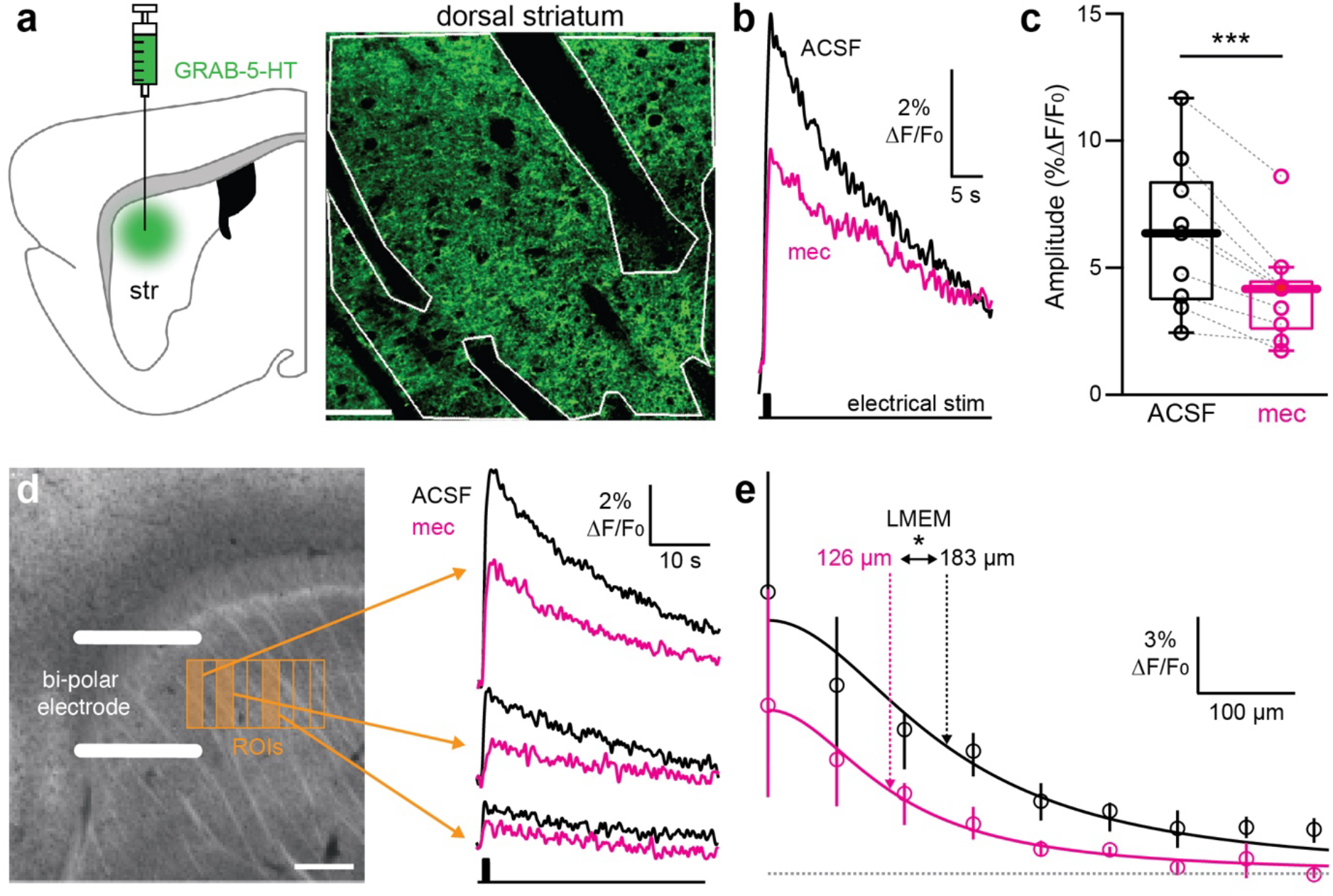
Nicotinic acetylcholine receptors (nAChRs) elevate and extend the spatial range of serotonin (5-HT) release evoked by intrastriatal electrical stimulation. (**a**) Schematic of adeno-associated virus (AAV) injection into the dorsolateral striatum (DLS) leading to GRAB-5HT expression (left). Representative field-of-view (FoV) of a region of DLS expressing GRAB-5HT imaged with two-photon laser scanning microscopy (2PLSM). White line indicates the region-of-interest (ROI). Scale bar 50 µm (right). (**b**) GRAB-5HT response to electrical stimulation (10 pulses at 10 Hz, 2 ms), before (black) and after (magenta) application of 10 µM mecamylamine (mec). (**c**) Distribution of peak GRAB-5HT responses before and after application of 10 µM mec. (**d**). Schematic of electrical stimulation of sagittal slices through DLS indicates the location of the sequence of 100 µm wide rectangular ROIs used to measure the peak GRAB-5HT response as function of distance from the stimulating electrode. (left). Representative traces of GRAB-5HT responses averaged over individual ROIs (0, 200, 500 µm), before (black) and after (magenta) application of 10 µM mec (right). (**e**) Lorentzian fit of maximal GRAB-5HT responses as a function of distance from the stimulating electrode (d), before (black) and after (magenta) application of 10 µM mec. Vertical lines indicate interquartile intervals, and circles represent the mean response for each band. * *P < 0*.*05*; *** *P < 0*.*001*

**Figure 2.**
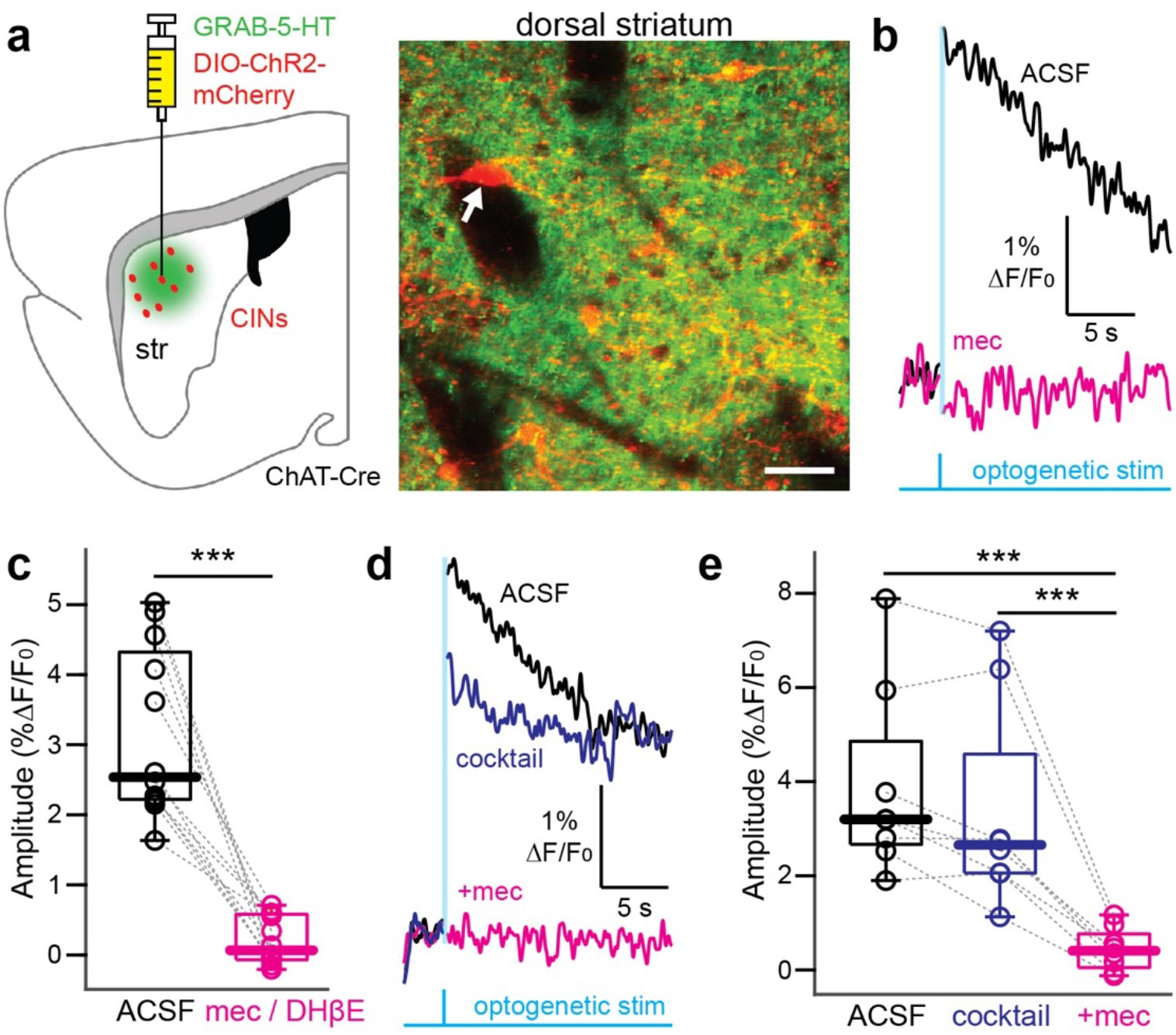
Optogenetic activation of striatal cholinergic interneurons (CIN) leads to nAChR-dependent 5-HT release in acute slices of the DLS. (**a**) Schematic of AAV injection into the DLS leading to non-selective expression of GRAB-5HT and selective expression of channelrhodopsin-2 (ChR2)-mCherry in CINs in ChAT-cre mice (left). 2PLSM image of a representative region of DLS expressing GRAB-5HT (green) overlaid with a Z-stacked image of CINs (white arrow) expressing ChR2-mCherry. Scale bar 50 µm (right). (**b**) GRAB-5HT response to a brief (2 ms long) optogenetic stimulation (470 nm), before (black) and after (magenta) application of 10 µM mecamylamine (mec). Vertical cyan line indicates a brief period of blanking of the stimulation artifact (see Methods). (**c**) Distribution of peak GRAB-5HT responses before (black) and after (magenta) application of nAChR blockers (10 µM mec or DHBE data pooled). (**d**) Same as (**b**), with the additional blue trace indicating the response after the application of a cocktail of glutamatergic, GABAergic and muscarinic receptor blockers (50 µM D-AP5, 10 µM DNQX, 2 µM CPG55845, 10 µM SR95531, and 10 µM atropine). (**e**) Same as (**c**) for the data in (**d**). *** *P < 0*.*001*.

We and others have shown that activation of nAChRs on nigrostriatal axon terminals not only increases the *amount* of DA release, but also nearly doubles the *spatial range* of its release^8,9^. We therefore measured the spatial extent of GRAB-5-HT responses by quantifying evoked fluorescence at 100 µm-wide increments from the stimulating electrode and fitting a Lorentzian function to the maximal responses (*N = 3* mice, *n = 9* slices) as a function of distance (**Fig. 1d,e**). The Lorentzian fell off with a length scale of 183 µm under control conditions. Mecamylamine application not only reduced the overall amplitude of the response, but also significantly shortened the length scale of the Lorentzian to 126 µm (*t*_*156*_ *= 2*.*583, P = 0*.*01*, LMEM), indicating that nAChRs increase the spatial extent of striatal 5-HT release by approximately 45% (**Fig. 1e**).

We observed that intrastriatal electrical stimulation promotes nAChR-sensitive release of 5-HT, but can endogenous ACh released from CINs drive nAChR-dependent release of striatal 5-HT on its own? To this end we inoculated the dorsolateral striatum of ChAT-Cre mice (which express Cre recombinase under the choline acetyltransferase promoter in CINs) with a mixture of the AAVs harboring GRAB-5-HT (broadly labeling the dorsal striatum) and DIO-channelrhodopsin-2 (ChR2)-mCherry (specifically labeling CINs) (see Methods) (**Fig. 2a; Supplementary Fig 2a**). Brief (2 ms duration) 470 nm LED stimulation resulted in a significant increase in GRAB-5HT fluorescence, with a slow decay lasting 10s of seconds (**Fig. 2b**). Thus, synchronous activation of CINs is sufficient to evoke striatal 5-HT release. The GRAB-5-HT signal was completely abolished by 10 μM mecamylamine or dihydro-β-erythroidine (DHβE), suggesting that CIN-evoked 5-HT release is mediated by α4β2 type nAChRs (*N = 4* mice, *n = 12* slices with mecamylamine, *n = 3* slices with DHβE *t*_*22*_ *= –9*.*061, P = 7*·*10*^*–9*^, LMEM) (**Fig. 2b,c**).

Is CIN control of 5-HT release direct, or are the key ACh receptors on striatal loci other than serotonergic axons? To rule out the involvement of postsynaptic nAChRs on GABAergic interneurons^21^, presynaptic nAChRs on glutamatergic inputs^22^ or muscarinic ACh receptors in general^23,24^ we repeated the above experiment in the presence of GABA, glutamate and muscarinic ACh receptor antagonists (see Methods). This antagonist cocktail had no significant effect on CIN-evoked 5-HT release, which was eliminated by subsequent blockade of nAChRs (**Fig 2d,e**). Together, these data suggest that ACh acts directly on nAChRs located on 5-HT fibers to mediate 5-HT release [*N = 3* mice, *n = 8* slices, *t*_*21*_ *= –7*.*322, P = 3*.*3*·*10*^*-7*^ (control *vs*. mec), *t*_*21*_ *= –5*.*454, P = 2*·*10*^*-5*^ (cocktail *vs*. mec) LMEM]. Furthermore, as the increase in GRAB-5-HT fluorescence induced by synchronous optogenetic activation of CINs is similar to the nAChR-dependent component of the GRAB-5-HT signal evoked by electrical stimulation (**Supplementary Fig. 3**) (*t*_*19*_ *= 0*.*962, P = 0*.*35*, LMEM), CINs are likely the sole source of ACh that promotes release of 5-HT. Nevertheless, to determine whether nAChRs amplify a putative co-release of 5-HT from DA axons, or any other mechanism by which excitation of striatal DA axons may indirectly cause the release of striatal 5-HT, we inoculated 1) the substantia nigra pars compacta (SNc) of TH-Cre mice (which express Cre-recombinase under the tyrosine hydroxylase promoter) with an AAV harboring DIO-ChR2-mCherry and 2) the dorsal striatum with an AAV harboring GRAB-5HT. Direct optogenetic activation of DA fibers induced a small but significant increase in GRAB-5HT. The increase, however, was only about a fifth of the size of the GRAB-5HT signal generated by synchronous optogenetic activation of CINs (**Fig. 2**) and was insensitive to mecamylamine, indicating that DA axons do not contribute to the nAChR-dependent release of striatal 5-HT (**Supplementary Fig. 4**).

**Figure 3.**
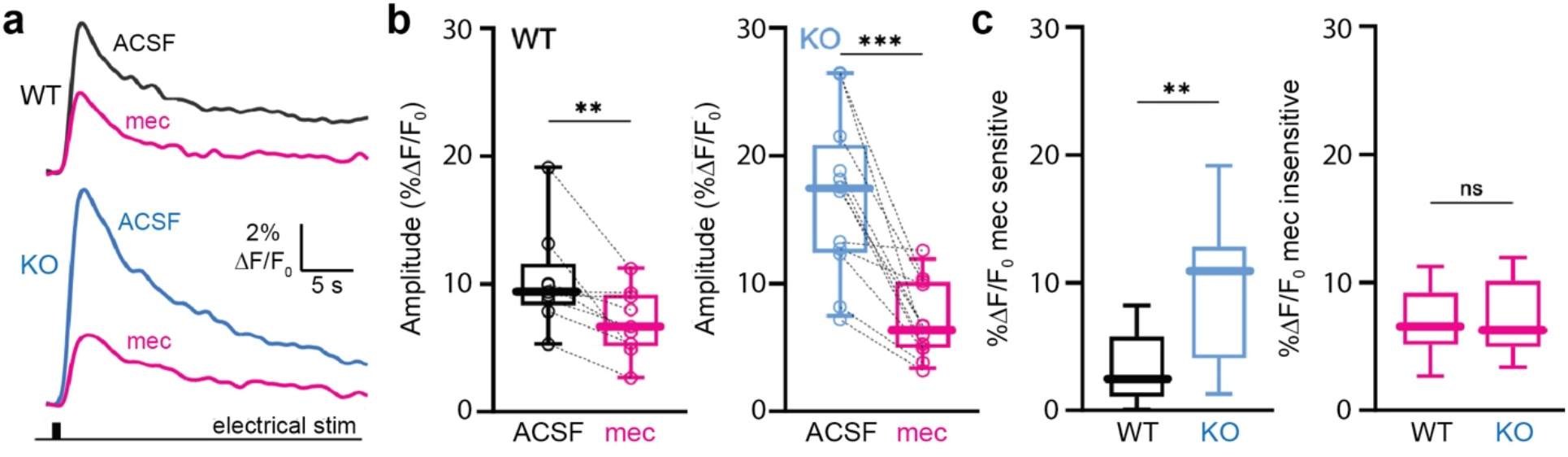
Increased cholinergic-mediated striatal serotonin release in Sapap3 KO mice. (**a**) Representative line scan traces of GRAB-5-HT fluorescence in the dorsolateral striatum of wild type (WT) (top, black) and knockout (KO) Sapap3 (bottom, blue) mice, before and after bath application of mecamylamine (mec, magenta). (**b**) Distribution of peak GRAB-5HT responses before (black – WT, cyan - KO) and after (magenta) application of mecamylamine in WT (left, *N = 5* mice, *n = 9* slices *t*_*16*_ *=* –3.5054, *P* = *0*.*002* LMEM) and KO (right, *N = 6* mice, *n = 12* slices *t*_*22*_ *= –6*.*066, P = 4*.*2· 10*^*-6*^ LMEM). (**c**) Quantification of mec-sensitive (left, *t*_*19*_ *= –2*.*890, P* = *0*.*009*) and -insensitive (right, *t*_*19*_ *= –9*.*5·* 10^−5^ *P = 0*.*999*, LMEM) components of GRAB-5HT signal in WT and KO mice. * *P < 0*.*05*; ** *P < 0*.*01;* *** *P < 0*.*001;* ns – not significant.

If CINs induce the release of 5-HT under physiological conditions, then in a pathophysiologically hypercholinergic striatum this mechanism should be amplified. We recently reported that Sapap3^-/-^ mice, which display OCD-like motor behaviors such as compulsive grooming, exhibit a hypercholinergic phenotype in the striatum: the density of CINs is higher, their autonomous firing rate is higher and they release more ACh in response to electrical stimulation in acute striatal slices^25^. We therefore asked if cholinergic modulation of 5-HT release is amplified in Sapap3^-/-^ mice. Indeed, intrastriatal electrical stimulation induced significantly larger GRAB-5-HT fluorescence signals in the dorsal striatum of Sapap3^-/-^ mice, and this difference disappeared in the presence of mecamylamine (**Fig. 3**) Supporting the conclusion that elevated 5-HT release is the result of increased CIN-mediated ACh release, only the mecamylamine-sensitive component of the GRAB-5-HT signal was elevated in Sapap3^-/-^ mice (**Fig. 3c**), suggesting that basal striatal 5-HT release capacity is unaltered. Corroborating that ACh facilitates 5-HT release by a mechanism similar to the one by which it facilitates DA release, intrastriatal electrical stimulation also induced significantly higher nAChR-sensitive (but not nAChR-insensitive) striatal DA release in the Sapap3^-/-^ mice (**Supplementary Fig. 5**).

Overall, this is consistent with the observation that 5-HT turnover (but not baseline levels) is elevated in Sapap3^-/-^ mice^26^.What does this imply for the etiology of OCD-like behaviors? The first line pharmacological treatment for OCD is selective serotonin reuptake inhibitors (SSRIs); while the mechanism of action is typically thought to involve increasing 5-HT tone, conclusive proof is lacking and some evidence suggests that chronic SSRI treatment may suppress 5-HT synthesis^27,28^. The answer to this question is beyond the scope of our current study, but our data offer a mechanism for monoamine dysregulation in a mouse model of compulsive behavior.

While the presence of nAChRs on 5-HT axons in the striatum was previously reported^15,16^, to the best of our knowledge, this is the first functional demonstration that activation of CINs can induce local 5-HT release, a phenomenon that extends our understanding of reciprocal acetylcholine-monoamine regulation to include 5-HT^8,9,11,14,19,20,29^. Specifically, as previously shown for to DA^8,9^, ACh released by CINs can directly induce, amplify and extend the spatial extent of striatal 5-HT release via activation of axonal nAChRs. This amplification is exaggerated in the striatum of Sapap3^-/-^ mice, which exhibit pathologically elevated CIN activity^25^, suggesting that nicotinic amplification of monoamine release may be a general property of hypercholinergic states in the striatum. Hypercholinergic states arise in the striatum in other neurological disorders, notably Parkinson’s disease (PD)^30^, raising the possibility that exaggerated recruitment of 5-HT release by CINs may contribute to PD pathophysiology.

Finally, given the growing evidence implicating CINs in levodopa induced dyskinesia (LID)^31,32^, it is tempting to speculate that abnormal CIN activity could amplify the “false” release of DA synthesized from levodopa in 5-HT terminals after dopamine denervation, leading to dyskinetic symptoms^33^.

## Acknowledgments

This work was funded by grants from the US-Israel Binational Science Foundation (no. 2021212) to J.LP. and J.A.G, from the Israel Science Foundation (no. 1959/22) to J.A.G and from NIH (NS 104089) and The Hartman Center to J.L.P. We would like to thank Prof. Ayal Ben-Zvi and Dr. Eran Lottem for valuable discussions; and Anatoly Shapochnikov and Drs. Yael Feinstein-Rotkopf and Tamar Licht and for excellent technical assistance.

## Methods

### Animals

Experimental procedures adhered to and received prior written approval from the Institutional Animal Care and Use Committees (IACUC) of the Hebrew University of Jerusalem (HUJI, protocol number MD-20-16113-3) and Stony Brook University (SBU). Experiments were carried out in the following mouse strains: 1. C57BL/6J mice (Strain #:000664; Jackson Laboratories, Bar Harbor, ME, USA); 2. ChAT-IRES-Cre (Δneo) transgenic mice (stock number 031661; Jackson Laboratories, Bar Harbor, ME, USA); 3. TH-Cre 1 (Strain #:008601; Jackson Laboratories, Bar Harbor, ME, USA, RRID:IMSR_JAX:008601); and 4. Sapap3 conditional knock-in mice (Malgady et al., 2023). All Sapap3^-/-^ mice lacked both copies of the Sapap3 gene. Two-to-seven-month-old C57BL/6J and transgenic mice were used for experiments. Sapap3^-/-^ mice were 3-7 months old, when they exhibited compulsive grooming. Sex was not considered as a factor in the design of the experiments, and so mice of both sexes were used and their results pooled. All mice were housed under a 12-h light/dark cycle with food and water ad libitum. Ambient temperatures and humidity were 22 ± 2°C and 50 ± 10%, respectively.

### Stereotaxic surgeries

At HUJI: Mice were deeply anesthetized with isoflurane in a non-rebreathing system (2.5% induction, 1–1.5% maintenance) and placed in a stereotaxic frame (model 940, Kopf Instruments, Tujunga, CA, USA). Temperature was maintained at 35 °C with a heating pad, antibiotic synthomycine eye ointment was applied to prevent corneal drying, and animals were hydrated with a bolus of injectable saline (5 ml/kg) mixed with an analgesic (5 mg/kg meloxicam). To calibrate specific injection coordinates, the distance between bregma and lambda bones was measured and stereotaxic placement of the mice was readjusted to achieve maximum accuracy. A small hole was bored into the skull with a micro drill bit and a glass pipette with AAVs attached to Nanoject III system (Drummond) was slowly inserted at the relevant coordinates under aseptic conditions. The coordinates of dorsolateral striatum (DLS) injection were: +0.5 mm anteroposterior (AP); +/-2.3 mm mediolateral (ML); −2.8 mm dorsoventral (DV), and for the SNc: -3.1 mm anteroposterior (AP); +/-1.2 mm mediolateral (ML); −4.2 mm dorsoventral (DV) relative to bregma using a flat skull position. To minimize backflow, solution was slowly injected (1nl/second) and the pipette was left in place for 5–7 min before slowly retracting it. For 5-HT sensor experiments, a total amount of 500 nl of an adeno-associated virus serotype 5 harboring the GRAB-5HT3.5 construct (AAV9hSyn-5HT3.5; 2.20 × 10^13^ vg/ml; WZ Biosciences Inc. Lot No. 20201210), recently renamed g5-HT2h^18^, was injected into DLS. For optogenetic stimulations of CINs a total amount of 650 nl adeno-associated virus cocktail of serotype 5 harboring the GRAB-5HT3.5 construct and ChR2 construct (AAV5/ Ef1a-DIO-hchR2(H134R)-mCherry; 4× 10^12^ vg/ml; UNC vector core; Lot# AV4313-2A) was injected into DLS of ChAT-IRES-Cre mice; for optogenetic stimulation of dopamine axons 500 nl of GRAB-5HT3.5 was injected into the DLS and 250 nl of the ChR2 construct in to the SNc of TH-Cre mice.

At SBU (experiments involving Sapap3^-/-^ mice): Mice were induced and maintained under anesthesia via inhalation of vaporized isoflurane (3% induction, 1–2% for maintenance). Prior to any surgical procedures, sedation was confirmed via toe pinch, then animals were placed in a stereotaxic apparatus, the scalp was shaved and disinfected with 70% ethanol and Betadine solution, and eyes were covered with protective gel (Puralube Ophthalmic Ointment, Dechra Veterinary Products). A 1μL Hamilton Neuros micro-volume syringe was used for unilateral DLS injections (0.5-62 mm AP; −2.2-3 mm ML; −2.8mm DV) and manipulated via a motorized stereotaxic injector (Quintessential Stereotaxic Injector QSI, Stoelting Co.) controlled by Angle Two software (Leica Biosystems). 500nL of AAV9-hSyn-5HT3.5 (≥1×10^13^vg/ml) or AAV9-hSyn-dLight1.1 (≥1×10^13^vg/ml) were injected over 5 min and a further 5 min was allowed for diffusion before slowly withdrawing the needle. All mice were administered analgesic (Meloxicam, 5 mg/kg, subcutaneous injection) immediately after surgery and monitored daily for 72h.

### Acute slice preparation

At HUJI: After prolonged incubation time (at least 3 weeks) inoculated mice were deeply anesthetized with ketamine (200 mg/kg)–xylazine (23.32 mg/kg) and perfused transcardially with ice-cold modified artificial cerebrospinal fluid (ACSF) bubbled with 95% O_2_–5% CO_2_, and containing (in mM): 2.5 KCl, 26 NaHCO_3_, 1.25 Na_2_HPO_4_, 0.5 CaCl_2_, 10 MgSO_4_, 0.4 ascorbic acid, 10 glucose, and 210 sucrose. The brain was removed and sagittal slices sectioned at a thickness of 275 µm were obtained in ice-cold modified ACSF. Slices were then submerged in ACSF, bubbled with 95% O_2_–5% CO_2_, and containing (in mM): 2.5 KCl, 126 NaCl, 26 NaHCO_3_, 1.25 Na_2_HPO_4_, 2 CaCl_2_, 2 MgSO_4_, and 10 glucose, and stored at room temperature for at least 1 h prior to recording.

At SBU (experiments involving Sapap3^-/-^ mice): 2-5 weeks after surgery, acute coronal slices (275 μm) containing DLS were obtained from mice following anesthetization with ketamine/xylazine (100mg/kg / 7mg/kg) and transcardial perfusion with ice-cold artificial cerebral spinal fluid (ACSF) containing (in mM): 124 NaCl, 3 KCl, 1 CaCl_2_, 1.5 MgCl_2_, 26 NaHCO_3_, 1 NaH_2_PO_4_, and 14 glucose. Slices were cut using a VT-1000 S vibratome (Leica Microsystems, Buffalo Grove, IL) and transferred to a holding chamber where they were incubated at 32°C for 45 min in ASCF containing (in mM) 2 CaCl_2_ and 1 MgCl_2_, then kept at room temperature until recording. During recordings, slices were perfused with ACSF heated to 32°C. All solutions were continuously bubbled with carbogen (95% O_2_ and 5% CO_2_).

### Slice visualization and 2PLSM imaging

At HUJI: Slices were transferred to the recording chamber of Femto2D-resonant scanner multiphoton system (Femtonics Ltd., Budapest, Hungary) and perfused with oxygenated ACSF at 32 °C. A ×16, 0.8 NA water immersion objective was used to examine the slice using oblique illumination. 2PLSM Ca^2+^/monoamine imaging: The 2PLSM excitation source was a Chameleon Vision 2 tunable pulsed laser system (680–1080 nm; Coherent Laser Group, Santa Clara, CA). Optical imaging of GRAB-5HT3.5 signals was performed by using a 920-nm excitation beam, cholinergic mCherry imaging was performed by using a 1060-nm excitation beam. The fluorescence emission was detected and collected with gated GaAsP photomultiplier tubes (PMTs) for high sensitivity detection of fluorescence photons as part of the Femto2D-resonant scanner. Field-of-view (FoV) of approximately 307 µm × 307 µm were selected and imaged at 31 Hz. Scans were performed using 0.6 μm pixels. Regions-of-interest (ROI) consisted of high fluorescent patches in the slice and were generated automatically via custom matlab code binarizing the image by pixel responses. The system is also equipped with full-field 470 nm LED illumination, which was used to stimulate the CINs in the ChAT-IRES-Cre mice transfected with DIO-hchR2(H132R)-mCherry. The gated PMTs are turned off for a time window of a few tens of milliseconds flanking the LED pulse. Nevertheless, there is a light artifact in the signal.

Electrical stimulation was carried out by a parallel bipolar Platinum-Iridium electrode with diameter of 125 µm and spacing of 500 µm (FHC, PBSA0575). The magnitude of the stimulus was controlled through a stimulus isolator (ISO-Flex, MicroProbes) while the frequency and duration were controlled by computer software (MESc, Femtonics). A total of 10 pulses (10 Hz, 2 ms duration, 3 mA) were delivered.

Time lapse imaging: One-second-long time lapse imaging (Supplementary Fig. 1) were taken with ∼30 seconds interval. Each point was calculated as the mean fluorescence intensity in the ROI (method explained above) over entire recording session (parameter mentioned above). 10µM dopamine or 10µM 5-HT were diluted in ACSF (see above) and perfused onto recording chamber for ∼ 15 minutes.

Optical and data were obtained using the software package MESc (Femtonics), which also integrates the control of all hardware units in the microscope & electrical stimulator. The software automates and synchronizes the imaging signals and stimulations (electric/ optical). Data was extracted from the MESc package (Femtonics) to personal computers using custom-made code in MATLAB (MathWorks, Natick, MA, USA) code. Fluorescent changes over time (Δ*F/F*_*0*_) datapoints were extracted such that Δ*F*/*F*_0_ =(*F* − *F*_0_)/*F*_0_, where *F* is the maximal fluorescent value recorded while evoking electrical / optogenetic stimulations; *F*_*0*_ denotes the averaged baseline fluorescence.

2PLSM resonant scanning: In Fig.1e, Δ*F/F*_*0*_ data points were fitted with a Lorentzian function *a*/[1+(*x*/*λ*)^2^] to extract the length scale λ of the fluorescence signals*’* decay with distance. Individual mice were considered independent samples. Visualized traces were smoothed using a gaussian filter (sigma = 150 ms for electrical stimulations, sigma = 60 ms for optogenetic stimulation).

At SBU (Slice 2PLSM 5-HT and dopamine sensor imaging in experiments involving Sapap3^-/-^ mice):

Recordings were made using an Ultima Laser Scanning Microscope system (Bruker Nano, Inc, Middleton, WI) equipped with an Olympus 60X/1.0 water-dipping lens (Olympus, Center Valley, PA) and Chameleon Ultra II laser (Coherent, Inc., Santa Clara, CA). To evoke 5-HT or dopamine release, a bipolar stimulating microelectrode (FHC Inc., Bowdoin, ME) powered by an ISO-Flex Stimulus Isolator (Microprobes for Life Science) was embedded into the surface of the slice just outside the 60x objective field-of-view (200μm^2^ FOV area). 2-photon laser line scans in the shape of a spiral (920nm, 10μs/pixel dwell, 21.2-83ms/line) covering the entire FOV were acquired while delivering electrical stimuli (1s at 10 Hz, 2 ms pulse width, 3 mA for 5-HT; single 1ms pulse for dopamine). Line scans began 2s before stimulation and continued for 57s (5-HT) or 8 ms (dopamine) after stimulation ended. 10µM mec or DHβE were then perfused onto the slice for 10 min and the line scan protocol was repeated. For baseline and post-drug recordings, three 60s recordings were averaged. *ΔF/F*_*0*_ was calculated using the average fluorescence of the first 1.5s of baseline for F_0_. The peak amplitude of the signal was defined as the maximal ΔF/F_0_ that occurred between 0-5s after stimulation.

### Immunohistochemistry and confocal imaging

After viral injection mice (see “stereotaxic surgeries”) mice were deeply anesthetized with ketamine-xylazine followed by cold perfusion to the heart of PBS and 4% PFA. The brain was removed and kept overnight at 4°C in 4% PFA. The next day, the brain was washed in PBS (three times, 20 min each) before thin 50 µm coronal slices were made using a vibratome (Leica VT1000S). Slices were briefly washed with CAS-BLOCK (Life Technologies) before being incubated in CAS-BLOCK (300 µl) overnight with Goat anti Vesicular-Acetylcholine-transporter antibodies (ABN100MI, Thermo Fisher, 1:200).The slices were then washed in PBS (three times, 20 min each) and incubated with CAS-BLOCK and secondary antibodies (Abcam ab6566, 1:200) for 3 hr. Antifade Mounting Medium (VECTASHIELD) was applied to prevent slice bleaching.

Confocal imaging were acquired by Spinning Disk confocal microscope (Nikon) using Yokogawa W1 Spinning Disk camera and a 50µm pinhole and a 20x dry objective with 0.75 N. A (CFI PLAN APO VC). Excitation laser used were 488 nm, 561nm, 638nm to excite GRAB-5HT, mCherry, anti VAChT respectively. Optical and data were obtained using the software package NIS (Nikon), which also integrates the control of all hardware units in the microscope. Data was extracted from the ND2 package to personal computers and visualized with Fiji software. Low-magnification images (**Supplementary Fig. 4**) were acquired using Nikon SMZ-25 fluorescent stereoscope equipped with ×1 and ×2 objectives.

### Statistical analysis

Due to the nested design of this experiment, wherein we sampled multiple slices from individual mice, we fit the data with a linear mixed-effects model (LMEM), where Δ*F/F*_*0*_ from each slice was modeled as a sum of a fixed effect product (e.g., drug vs. control) plus uncorrelated random effects due to each individual mouse. When fitting the Lorentzian the model was of a product of two fixed effect, which are the treatment and the distance, to fit the LMEMs to the data, we used the Matlab (Mathworks) FITLME command. We reported the number of slices, the number of mice, Student’s t_*dof*_ statistic (dof = degrees of freedom) and *P value*. The null hypotheses were rejected if the *P* value was below 0.05.

### Reagents used

Mecamylamine hydrochloride (mec), Sigma-Aldrich, Lot # 019M4108V CAS: 826-39-1; Atropine sulfate salt monohydrate, Sigma-Aldrich, Lot # BCBH8339V CAS No.: 5908-99-6; SR 95531 hydrobromide (Gabazine), Hello Bio, CAS: 104104-50-9; DNQX, TOCRIS, CAS: 2379-57-9; D-AP5, Hello Bio, CAS:79055-68-8; CGP 55845 hydrochloride, Hello Bio, CAS: 149184-22-5; Dihydro-β-erythroidine hydrobromide, TOCRIS, CAS: 29734-68-7; Citalopram hydrobromide(HB2142), Hello Bio, CAS [59729-32-7]; Dopamine hydrochloride (DA), Sigma-Aldrich, CAS No.: 62-31-7; RS 23597-190, TOCRIS, CAS No.: 149719-06-2. Serotonin (5HT), Sigma-Aldrich, CAS No.: 153-98-0.

## References

1. Balleine, B. W. & O’Doherty, J. P. Human and rodent homologies in action control: corticostriatal determinants of goal-directed and habitual action. Neuropsychopharmacology 35, 48–69 (2010).

2. Zhai, S., Tanimura, A., Graves, S. M., Shen, W. & Surmeier, D. J. Striatal synapses, circuits, and Parkinson’s disease. Curr. Opin. Neurobiol. 48, 9–16 (2018).

3. Mathur, B. N., Capik, N. A., Alvarez, V. A. & Lovinger, D. M. Serotonin induces long-term depression at corticostriatal synapses. J. Neurosci. 31, 7402–7411 (2011).

4. Pommer, S., Akamine, Y., Schiffmann, S. N., de Kerchove d’Exaerde, A. & Wickens, J. R. The Effect of Serotonin Receptor 5-HT1B on Lateral Inhibition between Spiny Projection Neurons in the Mouse Striatum. J. Neurosci. 41, 7831–7847 (2021).

5. Benloucif, S. & Galloway, M. P. Facilitation of dopamine release in vivo by serotonin agonists: studies with microdialysis. Eur. J. Pharmacol. 200, 1–8 (1991).

6. Mathur, B. N. & Lovinger, D. M. Endocannabinoid-dopamine interactions in striatal synaptic plasticity. Front. Pharmacol. 3, 66 (2012).

7. Surmeier, D. J., Carrillo-Reid, L. & Bargas, J. Dopaminergic modulation of striatal neurons, circuits and assemblies. Neuroscience 198, 3 (2011).

8. Liu, C. et al. An action potential initiation mechanism in distal axons for the control of dopamine release. Science 375, 1378–1385 (2022).

9. Matityahu, L. et al. Acetylcholine waves and dopamine release in the striatum. Nat. Commun. 14, 6852 (2023).

10. Kramer, P. F. et al. Synaptic-like axo-axonal transmission from striatal cholinergic interneurons onto dopaminergic fibers. Neuron 110, 2949–2960.e4 (2022).

11. Cachope, R. et al. Selective activation of cholinergic interneurons enhances accumbal phasic dopamine release: setting the tone for reward processing. Cell Rep. 2, 33–41 (2012).

12. Threlfell, S. et al. Striatal dopamine release is triggered by synchronized activity in cholinergic interneurons. Neuron 75, 58–64 (2012).

13. Blomeley, C. & Bracci, E. Excitatory effects of serotonin on rat striatal cholinergic interneurones. J. Physiol. 569, 715 (2005).

14. Virk, M. S. et al. Opposing roles for serotonin in cholinergic neurons of the ventral and dorsal striatum. Proc. Natl. Acad. Sci. U. S. A. 113, 734–739 (2016).

15. Schwartz, R. D., Lehmann, J. & Kellar, K. J. Presynaptic nicotinic cholinergic receptors labeled by [3H]acetylcholine on catecholamine and serotonin axons in brain. J. Neurochem. 42, 1495–1498 (1984).

16. Reuben, M. & Clarke, P. B. Nicotine-evoked [3H]5-hydroxytryptamine release from rat striatal synaptosomes. Neuropharmacology 39, 290–299 (2000).

17. Parent, M., Wallman, M.-J., Gagnon, D. & Parent, A. Serotonin innervation of basal ganglia in monkeys and humans. J. Chem. Neuroanat. 41, 256–265 (2011).

18. Deng, F. et al. Improved green and red GRAB sensors for monitoring spatiotemporal serotonin release in vivo. Nat. Methods 21, 692–702 (2024).

19. Zhou, F. M., Liang, Y. & Dani, J. A. Endogenous nicotinic cholinergic activity regulates dopamine release in the striatum. Nat. Neurosci. 4, 1224–1229 (2001).

20. Therelfell, C. Striatal dopamine release is triggered by synchronized activity in cholinergic interneurons - PubMed. https://pubmed.ncbi.nlm.nih.gov/22794260/.

21. Koós, T. & Tepper, J. M. Dual cholinergic control of fast-spiking interneurons in the neostriatum. J. Neurosci. 22, 529–535 (2002).

22. Lee, C.-H. & Hung, S.-Y. Physiologic Functions and Therapeutic Applications of α7 Nicotinic Acetylcholine Receptor in Brain Disorders. Pharmaceutics 15, 31 (2022).

23. Goldberg, J. A., Ding, J. B. & Surmeier, D. J. Muscarinic modulation of striatal function and circuitry. Handb. Exp. Pharmacol. 223–241 (2012) doi:10.1007/978-3-642-23274-9_10.

24. Abudukeyoumu, N., Hernandez-Flores, T., Garcia-Munoz, M. & Arbuthnott, G. W. Cholinergic modulation of striatal microcircuits. Eur. J. Neurosci. 49, 604–622 (2019).

25. Malgady, J. M. et al. Pathway-specific alterations in striatal excitability and cholinergic modulation in a SAPAP3 mouse model of compulsive motor behavior. Cell Rep. 42, 113384 (2023).

26. Wood, J., LaPalombara, Z. & Ahmari, S. E. Monoamine abnormalities in the SAPAP3 knockout model of obsessive-compulsive disorder-related behaviour. Philos. Trans. R. Soc. Lond. B. Biol. Sci. 373, 20170023 (2018).

27. Robbins, T. W., Vaghi, M. M. & Banca, P. Obsessive-Compulsive Disorder: Puzzles and Prospects. Neuron 102, 27–47 (2019).

28. Honig, G., Jongsma, M. E., van der Hart, M. C. G. & Tecott, L. H. Chronic citalopram administration causes a sustained suppression of serotonin synthesis in the mouse forebrain. PloS One 4, e6797 (2009).

29. Blomeley, C. & Bracci, E. Excitatory effects of serotonin on rat striatal cholinergic interneurones. J. Physiol. 569, 715–721 (2005).

30. Barbeau, A. The Pathogenesis of Parkinson’s Disease: A New Hypothesis. Can. Med. Assoc. J. 87, 802 (1962).

31. Choi, S. J. et al. Alterations in the intrinsic properties of striatal cholinergic interneurons after dopamine lesion and chronic L-DOPA. eLife 9, e56920 (2020).

32. Tubert, C., Paz, R. M., Stahl, A. M., Rela, L. & Murer, M. G. Striatal cholinergic interneuron pause response requires Kv1 channels, is absent in dyskinetic mice, and is restored by dopamine D5 receptor inverse agonism. eLife 13, (2024).

33. Sellnow, R. C. et al. Regulation of dopamine neurotransmission from serotonergic neurons by ectopic expression of the dopamine D2 autoreceptor blocks levodopa-induced dyskinesia. Acta Neuropathol. Commun. 7, 8 (2019).

