## Supplementary Figures for "Synchronous activation of striatal cholinergic interneurons induces local serotonin release"

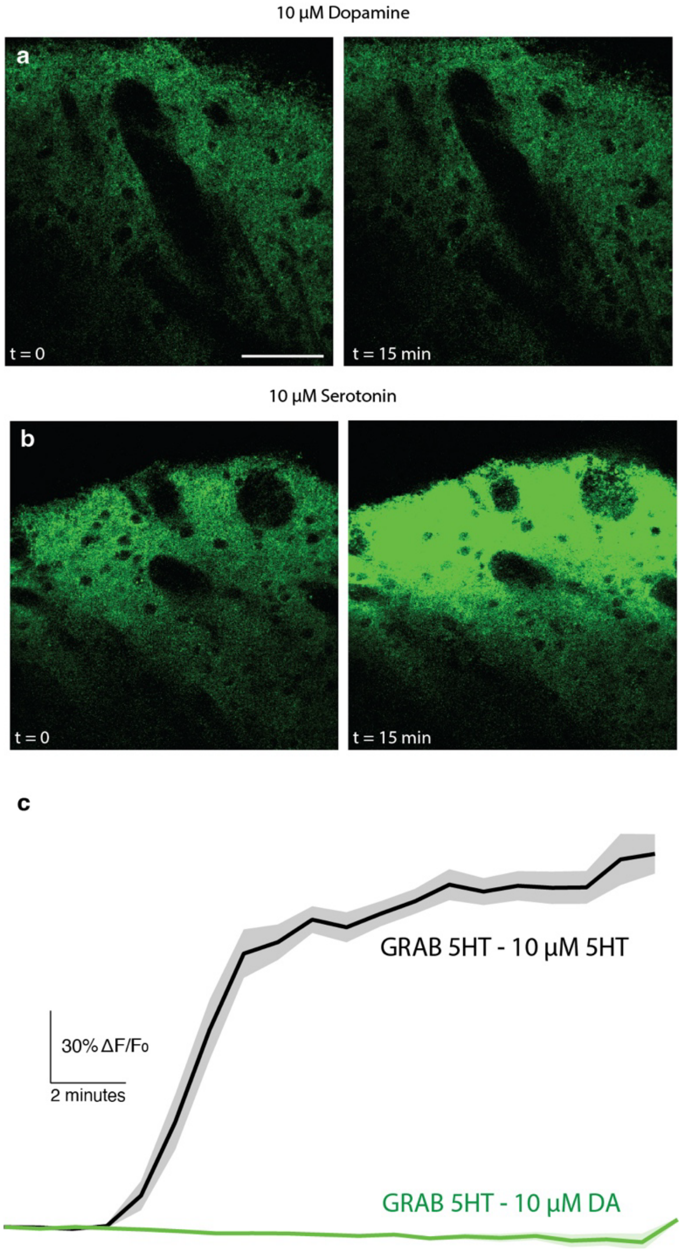


**Supplementary Figure 1. GRAB-5HT is insensitive to dopamine application.**
(**a**) Representative 2PLSM image of a region of DLS expressing GRAB-5HT in response to 10 μM dopamine at *t = 0* (left) and *t = 15* min (right). Scale bar 50 μm. (**b**) Same as (**a**), but in response to 10 μM serotonin. (**c**) Time lapse series of average fluorescence as function of time from (**a**) and (**b**).

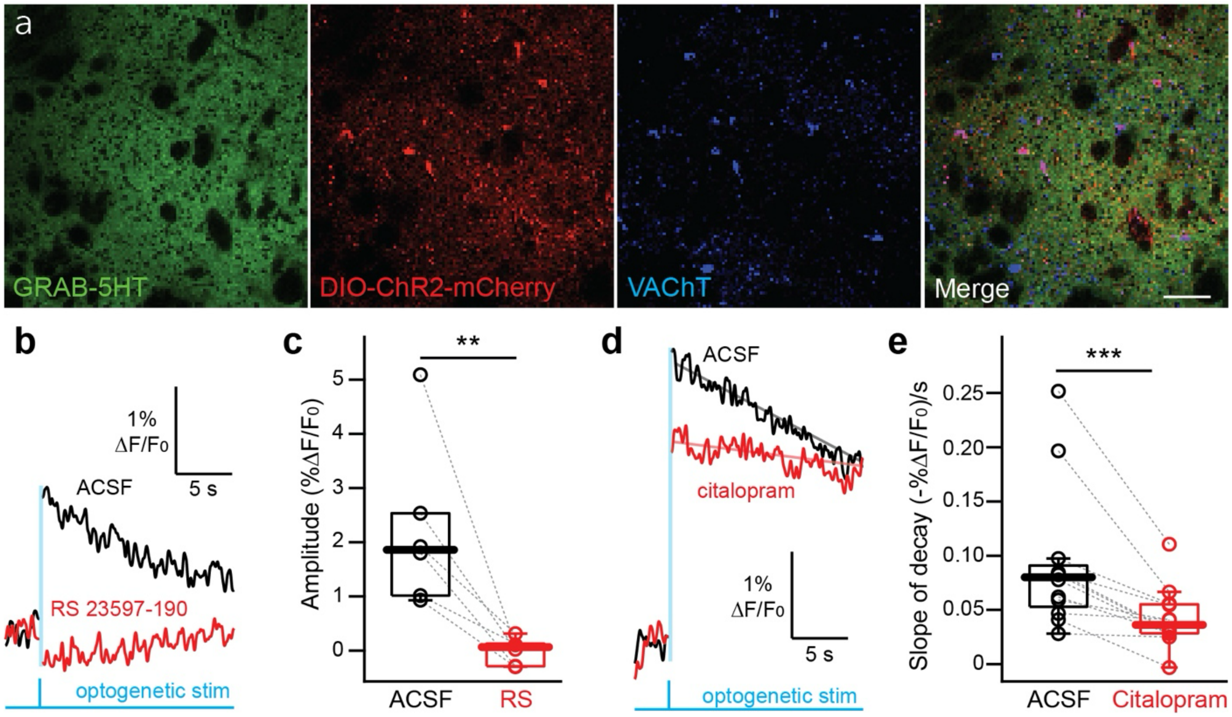


**Supplementary Figure 2. GRAB-5HT response to optogenetic activation of CINs is selective to 5-HT**.(**a**) Immunohistochemical staining of coronal striatal brain slices transfected with GRAB-5HT and DIO-ChR2-mCherry, along with anti-VAChT antibody, showing selective expression of ChR2 in ChAT neurons. Scale bar 100 µm (**b**) GRAB-5HT response to brief (2 ms) optogenetic stimulation (470 nm) before (black) and after (red) application of 35 µM RS 23597-190, a selective blocker of the 5-$HT_{2C}$ receptor on which GRAB-5HT is designed. (**c**) Distribution of peak GRAB-5HT responses to optogenetic stimulation, before (black) and after (red) RS 23597-190. (**d**) GRAB-5HT response to brief optogenetic stimulation before (black) and after (red) application of 30 µM citalopram, a selective 5-HT reuptake inhibitor, which slows decay kinetics. (**e**) Distribution of the slope of the decay of the GRAB-5HT trace before (black) and after (red) citalopram indicating a significant slowing of decay kinetics after citalopram. ** *P < 0.01*; *** *P < 0.001*

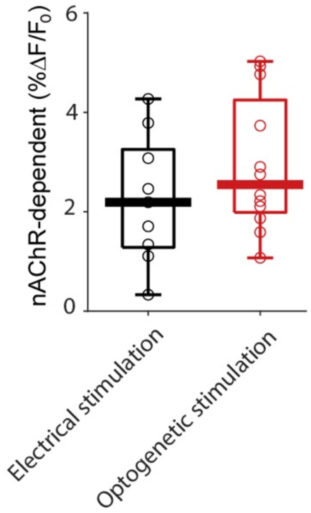


**Supplementary Figure 3. nAChR-dependent components of 5-HT release in DLS in response to electrical stimulation and optogenetic activation of CINs are equal**. Distributions of the nAChR-dependent component of the peak GRAB-5HT response (ACSF-mecamylamine) in the experiments with electrical stimulation (**Fig. 1c**) and optogenetic activation of CINs (**Fig. 2c**).

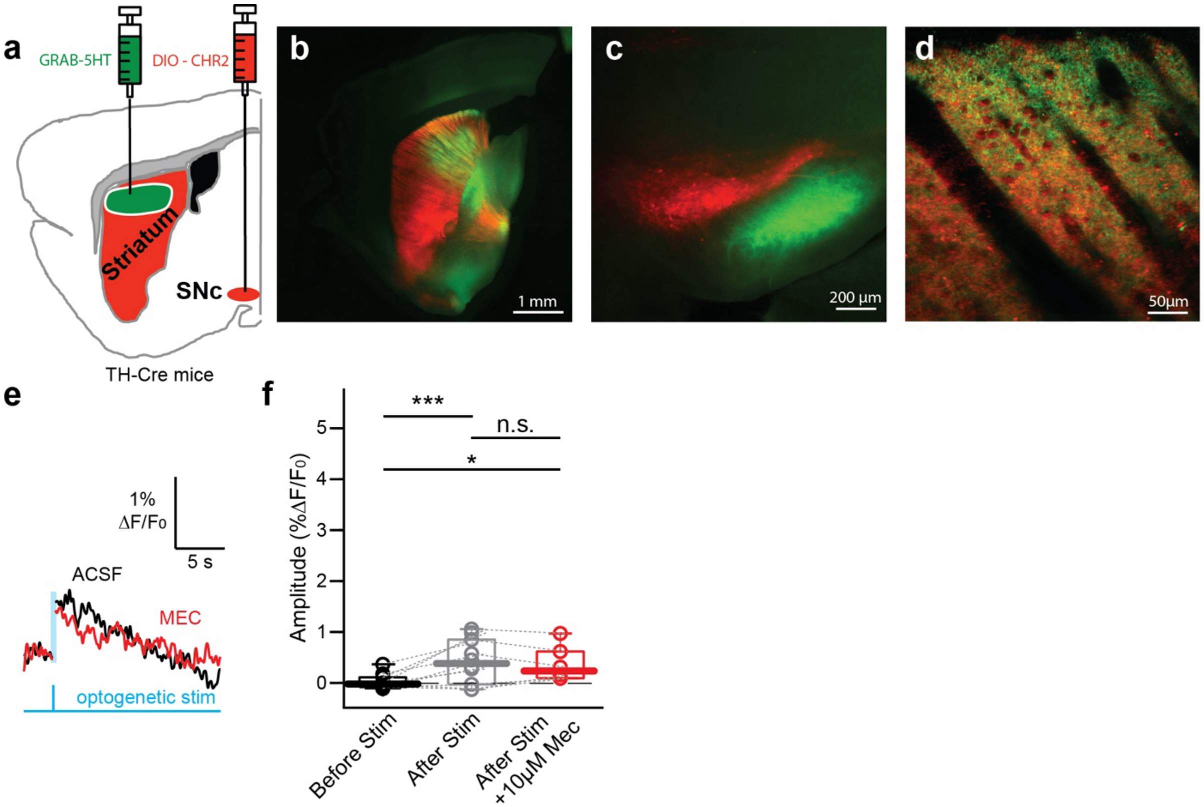


**Supplementary Figure 4. Optogenetic activation of striatal DA fibers elicits a small nAChR-independent GRAB-5HT response.** (**a**) Schematic of AAV injections in TH-Cre mice leading to non-selective expression of GRAB-5HT (green) in the DLS and selective expression of channelrhodopsin-2 (ChR2)-mCherry in dopaminergic neurons in the SNc (red). (**b**) sagittal slice through the striatum showing viral expression (**c**) Expression of viral vectors in the SNc (red) and striatal axons in the substantia nigra pars reticulata (green) in a coronal section of the midbrain. (**d**) 2PLSM image of a representative region of DLS expressing GRAB-5HT (green) overlaid with a Z-stacked image of DA neuropil expressing ChR2-mCherry. (**e**) Representative trace of GRAB-5HT response to a brief (2 ms long) optogenetic stimulation (470 nm) before (black) and after (red) application of 10 µM mecamylamine (Mec). (**f**) Distribution of baseline GRAB-5HT levels before optogenetic stimulation (black) and of peak responses before (grey) and after 10 µM Mec (red). The peak response was significantly higher than the baseline value ($N = 2$ mice, $n = 10$ slices, , $t_{22} = 4.51$, $P = 0.00017$, LMEM) but was not significantly (n.s.) different from the peak response after Mec ($n = 5$ slices, $t_{12} = -0.251$, $P = 0.81$, LMEM). The peak response after Mec remained significantly higher that the baseline value ($t_{12} = 2.41$, $P = 0.033$, LMEM). * $P < 0.05$; *** $P < 0.001$.

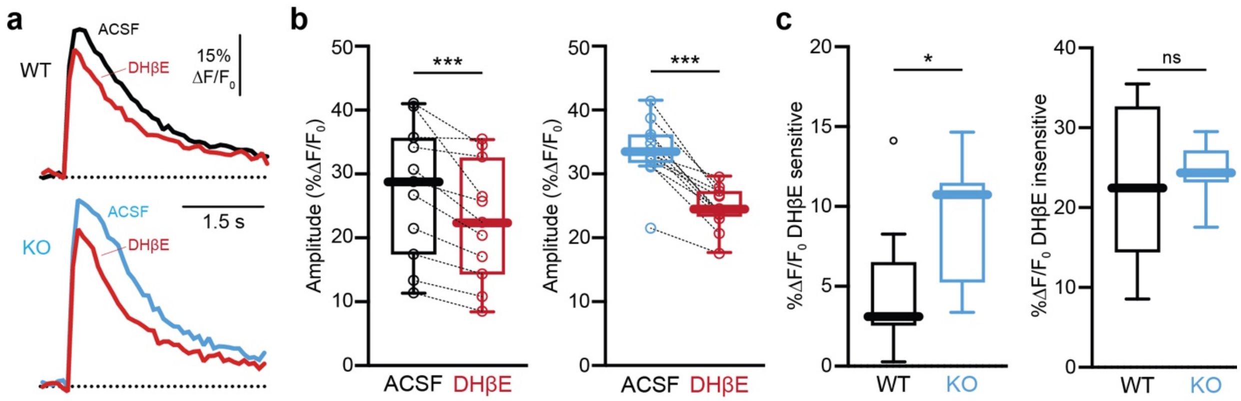


**Supplementary Figure 5. Elevated cholinergic tone contributes to increased DA release in Sapap3$^{-/-}$ mice.** (**a**) Averaged line scans measuring dLight1.1 fluorescence in dorsal striatal neuropil in WT (top, black) and KO (bottom, blue) mice before and after bath application of DHβE (red). (**b**) Distribution of peak dLight1.1 responses before (black) and after (red) in WT (black, left, $N = 4$ mice, $n = 11$ slices, $t_{20} = 4.322$, $P = 0.00033$, LMEM) and KO (blue, right, $N = 4$ mice, $n = 12$ slices, $t_{22} = 8.594$, $P = 1.76 \cdot 10^{-8}$, LMEM) mice. (**c**) Quantification of DHβE-sensitive ($t_{21} = 2.677$, $P = 0.014$, LMEM) and -insensitive ($t_{21} = -0.320$, $P = 0.752$, LMEM) components of peak dLight1.1 responses. * $P < 0.05$; *** $P < 0.001$; ns – not significant.
